# Fermentation-Induced Molecular Remodeling in African Indigenous Tubers: Cassava and Cocoyam

**DOI:** 10.64898/2026.06.05.730317

**Authors:** Alejandro Mendoza Cantu, Motseoa Mariam Lephatsi, Yusuf Abass Aleshinloye, Maserufe Faith Phahlane, Oluwaseun Peter Bamidele, Ntakadzeni Edwin Madala, Ndomele Ndiko Ludidi, Wout Bittremieux, Julia M. Gauglitz, Fidele Tugizimana

## Abstract

Cassava and cocoyam are major dietary staples in sub-Saharan Africa, commonly processed by natural fermentation before consumption. Although fermentation reduces antinutritional compounds and improves food quality, its molecular effects remain poorly characterized. We used untargeted mass spectrometry-based metabolomics with a computational annotation pipeline to compare fermentation-induced molecular remodeling in the two tubers, which showed distinct responses. In cassava, 718 of 773 significant features (92.9%) were depleted, indicating a predominantly catabolic process. In cocoyam, the response was more balanced, with 385 of 1,013 features (38.0%) enriched, including di- and tripeptides consistent with proteolytic processing. Class analysis, molecular networking, and pathway enrichment revealed tuber-specific signatures: cassava was dominated by purine metabolism, whereas cocoyam showed stronger enrichment of amino acid pathways. Cyanogenic glycoside-related features were depleted, consistent with detoxification. Biotransformation prediction also suggested putative fermentation products absent from current databases, highlighting the under-characterized chemistry of these tubers.

## 1. Introduction

Cassava (*Manihot esculenta*) and cocoyam (*Colocasia esculenta*) are important high-carbohydrate root crops, used for animal feeds, industrial applications, and most importantly, human consumption (Adebayo, 2023; Boakye et al., 2018). The starchy content of both plants has become the backbone of dietary energy in sub-Saharan Africa, shaping the local meals, food traditions, and economies. However, eating raw or undercooked cassava and cocoyam is both unpalatable and dangerous, as cassava contains toxic levels of cyanide, and cocoyam contains calcium oxalate crystals, both of which must be reduced through thorough processing. Particularly in cassava, fermentation has been the go-to method for addressing these antinutritional factors for generations, although the exact reasons why it works so well have been surprisingly underexplored. More recently, detailed biochemical studies of fermented cassava and cocoyam roots have shown that fermentation increases antioxidants, improves protein solubility, and reduces cyanide and mineral availability (Achidi et al., 2008; Bamidele, 2025).

Even with this growing biochemical evidence, our understanding is limited to metrics such as proximate composition, shifts in pH, titratable acidity, or microbial abundance, which do not inform us about the molecular scale. At the same time, the available tools have advanced dramatically. High-resolution mass spectrometry (MS) and newer computational metabolomics platforms now make it possible to track thousands of individual metabolites, even in foods as chemically diverse as fermented tubers. An untargeted MS study recently detected over 3,000 features across cassava genotypes, revealing far greater chemical diversity than previously recognized (Mantewu et al., 2025). Combined with emerging evidence of active metabolic turnover, peptide formation, and microbial community shifts during fermentation (Oguntoyinbo & Dodd, 2010; Yang et al., 2025), these findings suggest that a richer biomarker landscape has likely been overlooked. Together, they point to the need and opportunity for a metabolomics-driven study of fermentation-induced molecular remodeling.

Microbial fermentation is increasingly recognized as a biological process that actively reshapes the chemical composition of plant-based foods, generating diverse metabolites through the interplay of native enzymes, microbial communities, and endogenous phytochemicals (Sawant et al., 2025). Some of these changes improve nutrient accessibility or reduce antinutritional components, while others may produce molecules with physiological relevance that we are only beginning to understand. As global interest in fermentation science grows, African indigenous tubers present valuable opportunities for metabolomics studies that reveal how their biochemical profiles shift during fermentation.

Despite the availability of modern computational metabolomics tools, these techniques are rarely applied to African tubers. Cassava and cocoyam differ in starch structure, glycoside content, and phenolic composition, making it unlikely that they respond to microbial activity in the same way. By mapping the diversity of metabolites generated during fermentation and elucidating the biochemical pathways involved, we may begin to understand why fermented tubers are often safer, more palatable, and potentially more beneficial from a nutritional or functional perspective.

In this work, we apply a comparative untargeted metabolomics approach to map the molecular changes that fermentation induces in cassava and cocoyam, offering the first molecular-level view of how fermentation reshapes these culturally significant African tubers.

## 2. Materials and Methods

### 2.1 Extraction and preparation of samples

Cassava and cocoyam samples were purchased from local vendors in the markets of Thohoyandou, Limpopo province, South Africa. The harvested vegetables were transported to the pilot plant in the Department of Food Science and Technology at the University of Venda, washed, and processed into flour by milling after peeling the cassava and cocoyam tubers. The samples were processed according to standard protocols, including dicing into smaller pieces before milling. The milled samples were dried in an oven at 45 °C for 48 hours. The flour obtained was fermented for 72 hours, after which the fermentation was terminated by drying the samples in a hot air oven (Labotec EcoTherm 160L, Labotech, South Africa) at 40 °C for 12 hours. Unfermented samples served as controls. The samples were packed in zip-lock bags and stored at 8 °C. About 2 g of the dry and ground samples were weighed and extracted in 20 mL of 80% methanol by shaking on a digital rotisserie tube rotator (Dlab Scientific, Beijing, China) at 70 rpm, overnight. The crude extracts were centrifuged at 2739 g in a benchtop fixed-angle centrifuge (Thermo Fisher Scientific, Johannesburg, South Africa). The samples were then filtered through 0.22 µm nylon filters into high-performance liquid chromatography (HPLC) vials and kept at 4 °C until analysis by MS/MS. Romil SpS (Cambridge, UK) supplied LC-MS grade methanol, acetonitrile, and water. Analytical grade formic acid was supplied by Sigma-Aldrich (Johannesburg, South Africa).

### 2.2 LC-MS/MS data acquisition

Three biological replicates per condition (control and fermented) for each tuber were analyzed on a liquid chromatography-quadrupole time-of-flight (qTOF) MS/MS instrument (LCMS-9030 qTOF, Shimadzu Corporation, Japan). The chromatographic separation was performed on a Shim-pack Scepter C18-120 column (100 mm × 2.1 mm with particle size of 1.9 µm) (Shimadzu Corporation, Kyoto, Japan) kept at 55 °C. An injection sample volume of 3 µL was used with a binary mobile phase gradient which consisted of solvent A: 0.1% formic acid in Milli-Q water (both HPLC grade, Merck, Darmstadt, Germany) and solvent B: methanol (UHPLC grade, Romil SpS, Cambridge, UK) with 0.1% formic acid. A concave gradient elution was applied at a flow rate of 0.4 mL min⁻¹ over a total run time of 15 minutes. The gradient program consisted of 10% B for 2 min, a linear increase to 60% B over 3 min, followed by an increase to 90% B over 3 min, holding at 90% B for 3 min. The system was then returned to 60% B over 1 min and again to 10 % B in 1 min and finally the column was re-equilibrated for an additional 2 min prior to the next injection.

The chromatographic effluents were further analyzed utilizing the qTOF instrument set to acquire positive electrospray ionization data. The subsequent parameters were set as follows: interface voltage of 4.0 kV, interface temperature of 300 °C, nebulization and dry gas flow 3 L/min, heat block temperature of 400 °C, DL temperature of 280 °C, detector voltage of 1.8 kV, and the flight tube temperature at 42 °C. Sodium iodide (NaI) was used as a calibration solution to monitor high mass accuracy. MS1 and MS2 (through data dependent acquisition, DDA) were generated simultaneously for all ions with an *m*/*z* range between 100–1000 Da surpassing an intensity threshold of 2,000 counts. Fragmentation experiments were performed using argon as a collision gas at a collision energy of 30 eV with a spread of 5 eV.

### 2.3 Feature detection and alignment

Raw MS/MS data files were imported into MZmine (version 4.8.5) (Schmid et al., 2023) and processed separately for each tuber type. Mass detection was performed separately for MS1 and MS2 scans using an automated noise threshold, with minimum intensity cutoffs of 1,000 and 100 counts, respectively. Chromatogram building was restricted to a retention time window of 0.5 to 11.0 min to exclude the solvent front and late-eluting matrix signal. Extracted ion chromatograms were constructed requiring a minimum of four consecutive scans above 1,000 intensity and an absolute peak height threshold of 2,000 intensity, with a scan-to-scan *m*/*z* tolerance of 0.05 Da or 20 ppm. Prior to feature resolution, chromatograms were smoothed using a Savitzky-Golay filter over a window of five scans. Features were resolved using the local minimum resolver with a chromatographic threshold of 0.85, a minimum peak-to-edge intensity ratio of 1.8, and an allowed peak duration of up to 1.51 min. Isotopologue filtering was applied to retain only monoisotopic features using a tolerance of 0.001 Da / 3 ppm and a retention time window of 0.04 min, with a maximum considered charge state of 2. Features were aligned across all six samples of each tuber using the MZmine Join Aligner with tolerances of 0.05 Da / 8 ppm in *m*/*z* and 0.1 min in retention time, with *m*/*z* weighted three times more than retention time to prioritize mass accuracy. Missing values were recovered through gap filling within the same *m*/*z* and retention time windows, requiring a minimum of two data points. A duplicate feature filter was applied as a final cleanup step with tolerances of 0.0005 Da / 1.5 ppm and 0.035 min. Feature-level ion identity networking (Schmid et al., 2021) was performed to annotate adduct relationships, considering [M+H]+, [M+Na]+, [M+K]+, [M+NH4]+, and [M+H-H2O]+ ions.

### 2.4 Structural elucidation and chemical class assignment

Structural annotation was performed using SIRIUS (version 6.3.3) (Dührkop et al., 2019), with MS2 spectra exported directly from MZmine as input. Molecular formula prediction combined *de novo* fragmentation tree computation for features below *m*/*z* 400 and bottom-up search for features at or above *m*/*z* 400, with a fragment mass accuracy of 10 ppm. Structure candidates were retrieved by searching against biological databases available within SIRIUS, including FooDB (*FooDB*, n.d.), COCONUT (Chandrasekhar et al., 2025), and GNPS2 (Wang et al., 2016) spectral libraries. Features were considered high-confidence annotations when the SIRIUS confidence score reached 0.64 or above.

Chemical class ontology was assigned to all annotated features using CANOPUS (Dührkop et al., 2021), which maps molecular fingerprints to ClassyFire (Djoumbou Feunang et al., 2016) and NPClassifier (Kim et al., 2021) pathway classifications independent of database matching.

To expand the annotatable chemical space into fermentation-relevant transformation products, a sequential biotransformation strategy was integrated into the annotation pipeline. Predicted metabolites were generated using BioTransformer (version 3.0) (Wishart, Tian, et al., 2022), applying gut microbial and environmental microbial transformation rules across three iterative steps. In each step, the transformation products of the preceding step served as the substrate pool for the next, effectively simulating cumulative microbial biotransformation. The resulting predicted structures from all three steps were compiled and used in a second SIRIUS annotation pass, allowing tentative annotation of features that had remained unannotated in the initial search.

To assess representation across existing chemical databases, the SIRIUS database link field for each high-confidence annotation was parsed against ten reference databases: PubChem, ChEBI (Malik et al., 2026), LipidMaps (Fahy et al., 2009), HMDB (Wishart, Guo, et al., 2022), FooDB, COCONUT, NORMAN (Mohammed Taha et al., 2022), DSSTox (CCTE, 2025), SuperNatural (Gallo et al., 2023), and BioTransformer-predicted structures. Database intersection patterns were visualized as an UpSet plot with stacked bars indicating the contribution of each tuber.

### 2.5 Feature-based molecular networking

Feature-based molecular networking (Nothias et al., 2020) (FBMN) was performed on the GNPS2 platform using the aligned feature list and merged MS2 spectra exported from MZmine, independently for each tuber, with control and fermented conditions represented in a single network per tuber type. Two independent FBMN jobs were submitted: one searched against the standard GNPS2 spectral libraries, and a second incorporating the nearest neighbor suspect spectral library (Bittremieux et al., 2023) to extend annotation coverage toward structurally related but unannotated compounds. In both jobs, spectral library matching was performed using a cosine similarity threshold of 0.7, and edges between nodes in the molecular network were drawn based on a modified cosine similarity threshold of 0.7 with a minimum of five matched fragment ions. Network visualization and integration of annotation results from both jobs were performed in Cytoscape (version 3.10.3) (Shannon et al., 2003), where node attributes were merged into a consolidated annotated network for downstream interpretation.

### 2.6 Statistical analysis and differential abundance

The full feature tables exported from MZmine were processed in Python (version 3.12) using pandas (version 2.3.3) (McKinney, 2010; The pandas development team, 2025) and NumPy (version 2.1.3) (Harris et al., 2020). Peak areas were first normalized by sample weight to account for differences in extracted material across conditions. A global prevalence filter was then applied to retain only features detected in at least two out of three replicates in at least one of the four sample groups, ensuring a consistent feature set across all comparisons. Missing values remaining after filtering were imputed by replacing zeros with random values drawn from a uniform distribution between zero and the second lowest positive intensity observed across all samples, with a fixed random seed for reproducibility. Principal component analysis (PCA) was performed on the full log₂-transformed and weight-normalized feature matrix using PCA and StandardScaler from scikit-learn (version 1.7.2) (Pedregosa et al., 2011).

Differential abundance between fermented and control conditions was assessed separately for cassava and cocoyam using Welch’s t-test as implemented in SciPy (version 1.16.3) (Virtanen et al., 2020). Log₂ fold change (log₂FC) was calculated as the difference between the mean log₂-transformed intensities of the fermented and control groups. For volcano plot visualization, p-values were corrected for multiple testing using the Benjamini-Hochberg false discovery rate (FDR) procedure, with significance defined as an FDR-adjusted p-value below 0.05 and an absolute log₂FC greater than 1. Uncorrected p-values were retained as input for pathway analysis to preserve sensitivity for downstream enrichment testing. Pathway enrichment analysis was conducted using the Mummichog (Li et al., 2013) algorithm implemented in MetaboAnalyst (version 6.0) (Pang et al., 2024), with feature *m*/*z* values and uncorrected Welch’s t-test p-values as input. Analysis was run independently for cassava and cocoyam using the metabolic models of four microorganisms commonly associated with natural tuber fermentation: *Acetobacter*, *Lacticaseibacillus rhamnosus*, *Lactobacillus plantarum*, and *Saccharomyces cerevisiae*, complemented by lipid and non-lipid chemical reference models. Pathway significance was assessed internally by Mummichog using Fisher’s exact test, and pathways with a -log10(p-value) above 1 and at least two significant hits were considered enriched. The enrichment factor, calculated as the ratio of observed to expected hits, was reported to characterize the magnitude of enrichment. Results were compared across tuber types to identify shared and tuber-specific metabolic responses to fermentation.

To assess whether specific chemical classes were disproportionately affected by fermentation relative to the overall metabolome response, a two-sided Mann-Whitney U test was applied at each NPClassifier classification level (pathway, superclass, and class). For each class, the log₂FC values of features assigned to that class (CANOPUS probability ≥ 0.7) were compared against the log₂FC values of all remaining features. P-values were corrected for multiple testing using the Benjamini-Hochberg procedure within each classification level, adapted from the class-level significance approach described by Dührkop et al (Dührkop et al., 2021).

## 3. Results and Interpretation

### 3.1 Cassava and cocoyam show distinct metabolomic responses to fermentation

After feature detection and preprocessing, the feature tables contained 3,466 features for cassava and 4,468 features for cocoyam. PCA revealed clear separation between control and fermented samples in both tubers (**Figure 1a,b**). In cassava, the fermented samples cluster tightly while control samples are more dispersed, suggesting a smaller chemical variability after fermentation. In cocoyam the opposite was observed, with control samples clustering closely and fermented samples showing greater dispersion, pointing to a more diverse set of chemical changes introduced during fermentation.

**Figure 1.**
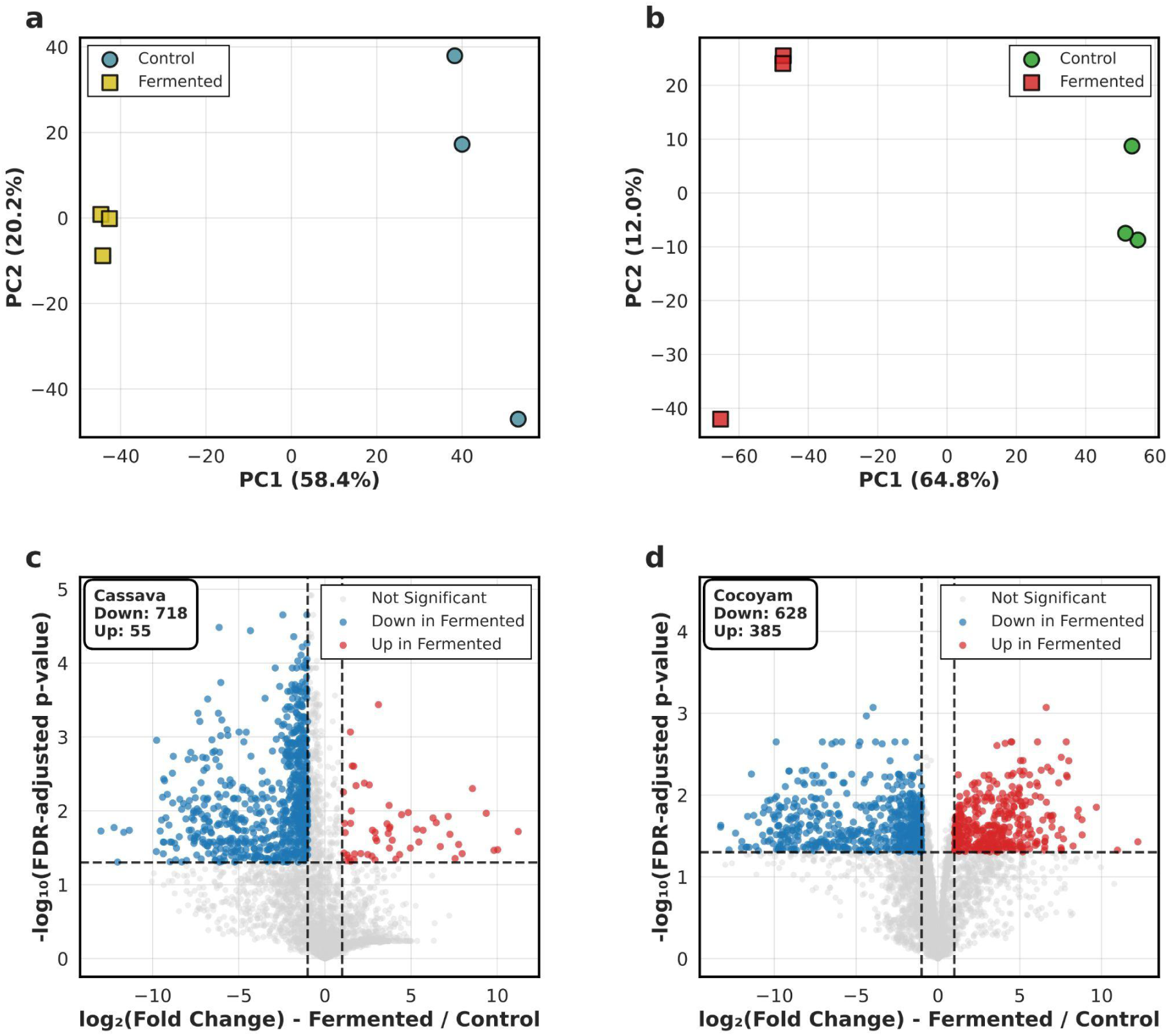
Cassava and cocoyam fermentations show different metabolomic profiles. **(a,b)** PCA score plots for cassava **(a)** and cocoyam **(b)** show separation between control and fermented conditions. **(c,d)** Volcano plots show differential abundance between control and fermented conditions for cassava **(c)** and cocoyam **(d)**, with significantly depleted features shown in blue and enriched features in red (FDR-adjusted p-value < 0.05, |log₂FC| > 1).

Differential abundance analysis confirmed these patterns. Fermentation in cassava displays a clear catabolic profile. Of the 773 significantly changed features, 718 (92.9%) were depleted during fermentation, with only 55 (7.1%) showing significantly higher abundance in the fermented samples (**Figure 1c**). This net depletion of features aligns with the well-documented reduction of starch and carbohydrate fractions during fermentation, which also affects physicochemical properties such as moisture content (Oyeyinka et al., 2020). In cocoyam, fermentation both depleted existing metabolites and generated detectable products, including smaller molecules likely derived from the breakdown of larger substrates. Of the 1,013 significant features found in cocoyam, 385 (38.0%) were enriched by fermentation while 628 (62.0%) were depleted (**Figure 1d**). These insights are consistent with reported increases in crude protein content and water absorption capacity in fermented cocoyam, reflecting a more active metabolic process (Oke & Bolarinwa, 2012).

### 3.2 Fermentation impacts distinct chemical classes in cassava and cocoyam

At the pathway level, the broadest NPClassifier level, most chemical classes in cassava showed reduced intensities in fermented samples compared to controls (**Figure 2a**). Fatty acids were the only pathway significantly depleted relative to the overall metabolome response (adj p = 0.004, n = 263), though visible decreases were also observed for amino acids, peptides, and carbohydrates. In cocoyam, no major shifts were observed at the pathway level (**Figure 2b**), with one exception: amino acids and peptides were significantly different from the background metabolome response (adj p = 0.0008, n = 190), though their near-zero median fold change reflects opposing trends within the class rather than a uniform shift.

**Figure 2.**
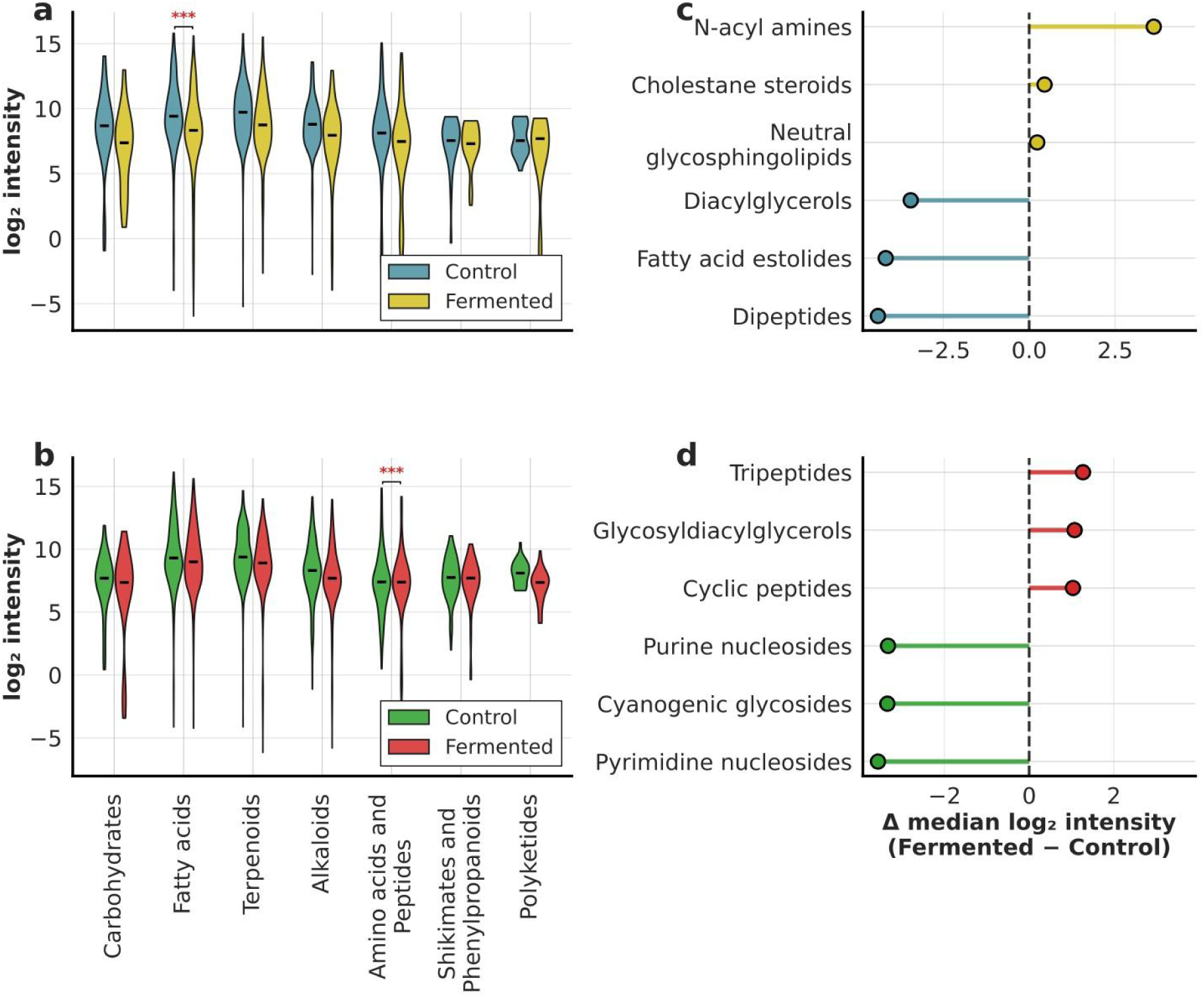
Chemical class-level intensity distributions and fermentation-induced shifts in cassava and cocoyam. **(a,b)** Violin plots show log₂-normalized intensity distributions of SIRIUS-annotated features grouped by NPClassifier pathway for **(a)** cassava and **(b)** cocoyam, comparing control and fermented conditions. Significance was assessed using a two-sided Mann-Whitney U test comparing log₂FC values of features within each class against all other features, with Benjamini Hochberg FDR correction (fatty acids in cassava: p = 0.004313; amino acids and peptides in cocoyam: p = 0.000759). Only features with NPClassifier pathway probability ≥ 0.7 and at least one real measured value in the respective tuber are included. **(c,d)** Lollipop plots show the top three most enriched and depleted NPClassifier classes upon fermentation for **(c)** cassava and **(d)** cocoyam, ranked by median log₂ intensity difference between fermented and control conditions.

Among the few classes enriched in fermented cassava samples, N-acyl amines are particularly notable (**Figure 2c**). These compounds, composed of amino acid and lipid moieties, have been investigated as microbial chemical signaling molecules (Mannochio-Russo et al., 2025). Their enrichment alongside glycosyldiacylglycerols, which are commonly found in the membranes of gram-positive bacteria (Blanc et al., 2013), suggests that detection in fermented cassava may partly reflect microbial biomass rather than transformations of the tuber metabolome itself.

At finer NPClassifier class levels, cocoyam showed more differentiation (**Figure 2d**). Neutral glycosphingolipids show increased abundance in fermented cocoyam, which may reflect membrane lipid remodeling associated with microbial or fungal activity during fermentation (Barreto-Bergter et al., 2006). In cassava, glycerolipids and glycerophospholipids were significantly depleted, while in cocoyam, small peptides were significantly enriched and fatty amides depleted (**Supplementary Figure 1a,b**). ClassyFire analysis at the superclass level further supported these patterns (**Supplementary Figure 2**). In cassava, only one feature classified by CANOPUS as a cyanogenic glycoside (*m*/*z* 490.19, [M+Na]+) was detected, and it was significantly depleted after fermentation (log₂FC = −6.67, **Supplementary Figure 3a**). In cocoyam, multiple features were classified as cyanogenic glycosides, with the class-level median showing a trend toward depletion (**Figure 2d**). The same feature found in cassava was also matched to a cyanogenic glycoside in cocoyam, but it did not show a significant reduction (**Supplementary Figure 3b**).

### 3.3 Fermented tubers have molecular diversity underrepresented in existing databases

Structural annotation was performed independently for the cassava and cocoyam datasets using SIRIUS and GNPS2 spectral library matching. At the feature level, 93 high-confidence annotations were obtained for cassava and 125 for cocoyam. These 218 annotated features correspond to 179 unique compounds by planar InChIKey, of which 51 were unique to cassava and 91 to cocoyam, with 37 shared between both tubers (**Figure 3a**). Molecular fingerprints were matched against ten reference databases (**Figure 3b**). Food-specific databases showed lower representation than general chemistry databases, with FooDB matching 99 of 218 features. In cassava, BioTransformer generated two putative structures not matched by any reference database, suggesting fermentation products that have not yet been cataloged.

**Figure 3.**
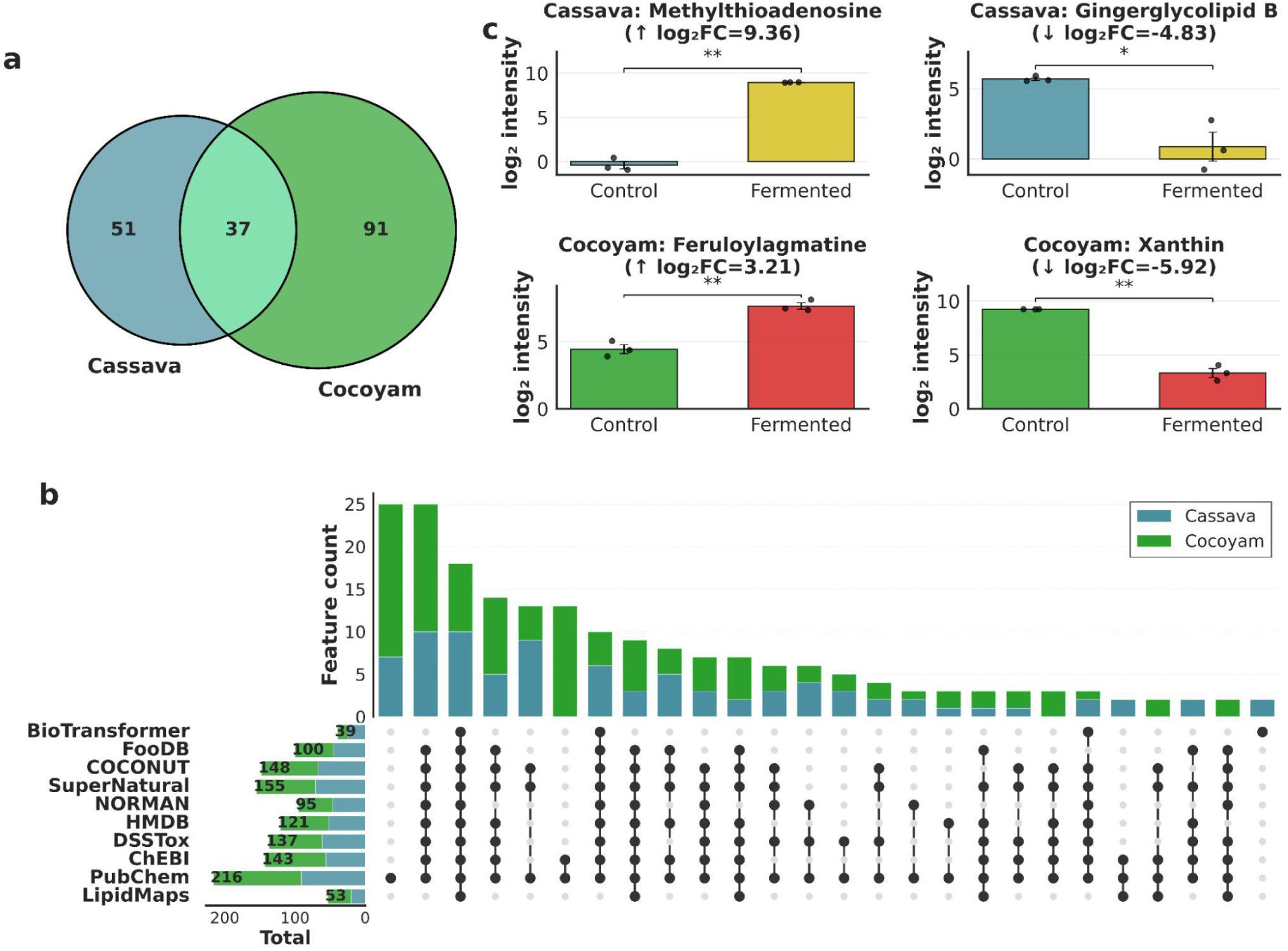
(a) Venn diagram showing the overlap of unique InChIKeys (GNPS + high-confidence SIRIUS combined) between cassava (51 unique) and cocoyam (91 unique), with 37 compounds shared between both tubers. **(b)** UpSet plot showing database intersection coverage of 218 high-confidence SIRIUS annotations (93 cassava, 125 cocoyam) across ten reference databases, with stacked bars indicating the contribution of each tuber. **(c)** Bar plots show log₂-normalized mean intensities (± standard error of the mean) of four representative annotated features comparing control and fermented conditions in cassava (top row) and cocoyam (bottom row). Individual replicates are shown as black dots. Significance brackets indicate Welch’s t-test results (* p < 0.05, ** p < 0.01).

Four compounds with significant differential abundance illustrate the molecular diversity in these matrices (**Figure 3c**). Two of these represent possible fermentation products identified using BioTransformer: methylthioadenosine and feruloylagmatine. Methylthioadenosine was significantly enriched in fermented cassava (log₂FC = 9.36, p < 0.01). This compound has been reported as a product of spermidine and spermine synthesis in *S. cerevisiae*, where over 98% of it originates from decarboxylated S-adenosylmethionine during polyamine biosynthesis (Chattopadhyay et al., 2006). It has also been identified as a metabolite produced by probiotic bacteria (Lyu et al., 2024). Its presence alongside two strongly enriched N-acyl amines (log₂FC = 10.03 and 9.81) reinforces the observation that the few enriched classes in fermented cassava reflect microbial activity rather than food-derived chemistry. In cocoyam, feruloylagmatine was enriched after fermentation (log₂FC = 3.21, p < 0.01; **Figure 3c**), consistent with its known role as a plant defense compound that accumulates under pathogen challenge and stress conditions (Muroi et al., 2009). Gingerglycolipid B, a monoacyldigalactosylglycerol originally described in ginger rhizome (Yoshikawa et al., 1994), was depleted in fermented cassava (log₂FC = −4.83, p < 0.05; **Figure 3c**), consistent with consumption of endogenous plant glycolipids during fermentation. Xanthine was depleted in fermented cocoyam (log₂FC = −5.92, p < 0.01) and as a purine degradation intermediate its reduction aligns with lactic acid bacteria activity reported in traditional fermented foods (Liu et al., 2025).

### 3.4 Molecular networking links structurally related fermentation products and substrates

The molecular networks of both tubers are dominated by sphingolipid and glycerolipid clusters, but differ in their diversity and fermentation response.

In cassava, the most densely connected regions contain monoacylglycerols and sphingolipids (**Figure 4a**), consistent with the reported lipid content of cassava involved in signaling and membrane protection (Mantewu et al., 2025). Terpenoid clusters are also present throughout the network. Depleted features are distributed broadly across the network with very few enriched nodes, consistent with the catabolic fermentation profile observed in the differential abundance analysis.

**Figure 4.**
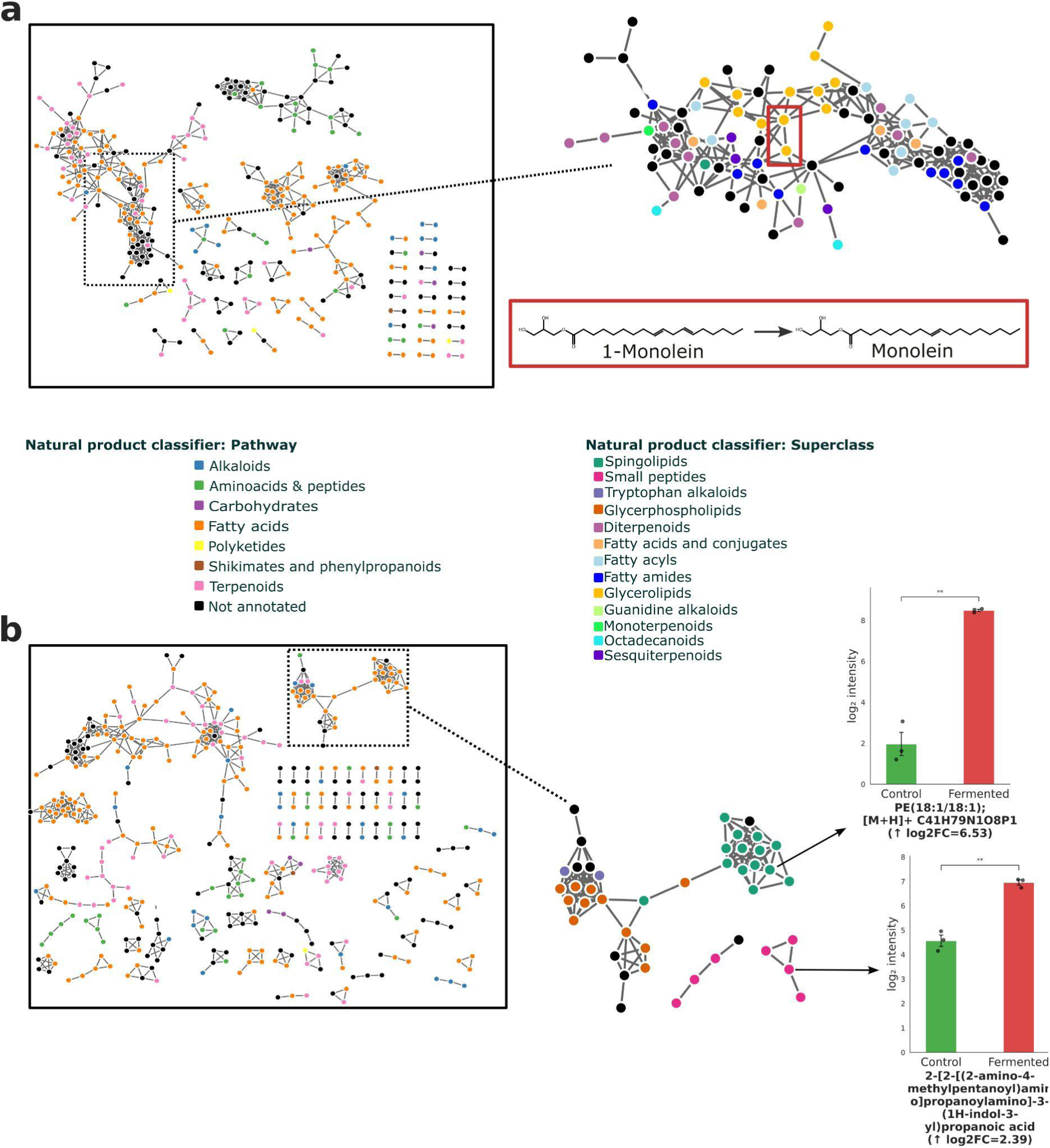
Feature-based molecular networking of cassava **(a)** and cocoyam **(b)**. Complete networks are shown on the left, colored by NPClassifier pathway classification. Selected subnetworks are expanded on the right, colored by NPClassifier superclass. **(a)** In cassava, the red box highlights a BioTransformer-predicted transformation between 1-monolinolein and monolein, confirmed by spectral similarity in the network. **(b)** In cocoyam, bar plots show two representative enriched features: a phosphatidylethanolamine (PE 18:1/18:1, log₂FC = 6.53, p < 0.01) and a tryptophan-containing tripeptide (log₂FC = 2.39, p < 0.01).

In cocoyam, a wider variety of compound families are connected across the network (**Figure 4b**). Enriched nodes tend to cluster together within specific molecular families, indicating that fermentation generates structurally related groups of new compounds rather than isolated metabolites. A phosphatidylethanolamine (PE 18:1/18:1, log₂FC = 6.53, p < 0.01) and a tryptophan-containing tripeptide (log₂FC = 2.39, p < 0.01) illustrate structurally distinct fermentation products found within their respective molecular families.

The BioTransformer annotation strategy provided an additional layer of interpretation. In cassava, 21 BioTransformer-predicted pairs were found within the same network component and three pairs were directly connected by spectral similarity edges, corresponding to a monoacylglycerol transformation chain (1-monolinolein, monoolein, glyceryl monostearate) and a sphingolipid series linking a neutral glycosphingolipid to three ceramide metabolites (**Figure 4a**). All of these features are depleted or not significant in the fermented condition, and none of the 37 BioTransformer-annotated features in cassava were enriched after fermentation. The overlap between predicted structural relationships and spectral network connections provides confidence that BioTransformer captures real substrate–metabolite pairs in the tuber.

In cocoyam, no BioTransformer pairs were directly connected by spectral edges, but 9 of 47 annotated features were enriched after fermentation, in contrast with cassava where this number was zero. Seven of these correspond to dipeptides and tripeptides, including Phe-Leu, Phe-Phe, and Leu-Ala-Phe, pointing to active proteolytic activity during cocoyam fermentation consistent with lactic acid bacteria acting on plant proteins (Ter et al., 2024).

### 3.5 Pathway enrichment reveals tuber-specific metabolic routes during fermentation

Pathway enrichment analysis was performed using metabolic models of four microorganisms commonly found in natural tuber fermentation (*Acetobacter, Lacticaseibacillus rhamnosus, Lactobacillus plantarum, and Saccharomyces cerevisiae*), complemented by lipid and non-lipid chemical subclass models. Most enriched pathways were driven by depleted features, mapping substrate consumption rather than product formation.

In cassava, purine metabolism was the clearest signal, enriched across all four microbial models (p = 0.0015 to 0.0078), with all 14 significant compound hits depleted in the fermented condition and none enriched (**Figure 5a**). Deoxyadenosine and deoxyguanosine were both significantly reduced, and purine salvage in *S. cerevisiae* is tightly linked to methionine recycling through methylthioadenosine (Thomas & Surdin-Kerjan, 1997), consistent with the enrichment of this compound observed above (**Figure 3c**). Folate biosynthesis appeared in one model (*Acetobacter*) and may reflect downstream consequences of active purine turnover.

**Figure 5.**
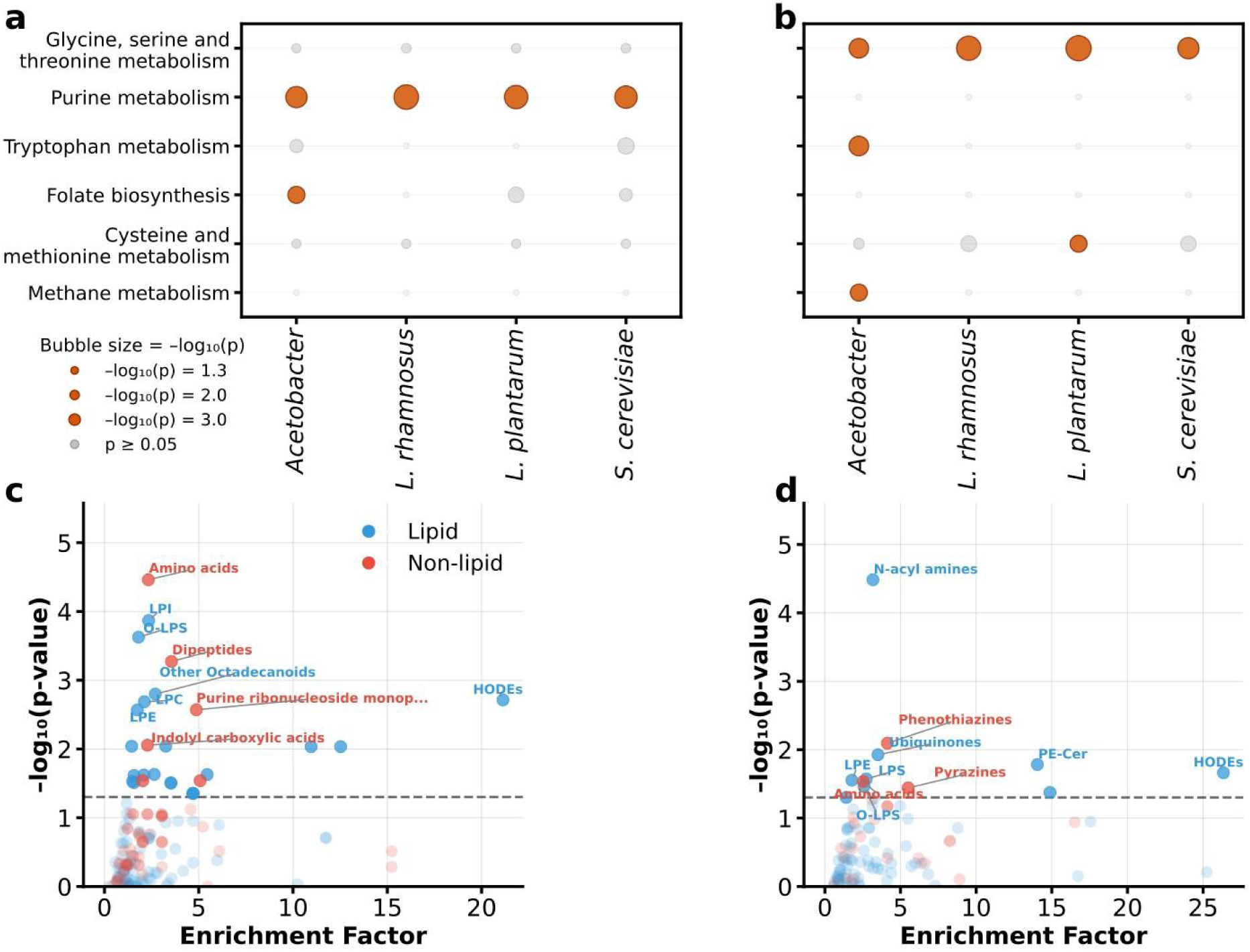
Enrichment analysis reveals tuber-specific metabolic routes activated during fermentation. **(a,b)** Bubble plot showing pathway enrichment results across four microbial metabolic models (*Acetobacter, L. rhamnosus, L. plantarum, S. cerevisiae*) for cassava **(a)** and cocoyam **(b)**, where bubble size represents -log₁₀(p-value) and colored bubbles indicate significance (p < 0.05). **(c,d)** Chemical subclass enrichment scatter plots show the enrichment factor versus -log₁₀(p-value) for combined lipid and non-lipid reference models in cassava **(c)** and cocoyam **(d)**. Labeled classes represent the most significantly enriched chemical subclasses in each tuber. The dashed line indicates p = 0.05.

In cocoyam, glycine, serine, and threonine metabolism was the dominant pathway, enriched across all four microbial models (p = 0.0011 to 0.0163) with enrichment ratios between 6 and 9 (**Figure 5b**).Tryptophan metabolism, cysteine and methionine metabolism, and methane metabolism were each enriched in one model (*Acetobacter*, *L. plantarum*, and *Acetobacter*, respectively). This amino acid-centered enrichment in cocoyam connects directly to the dipeptide and tripeptide generation observed in the class analysis (**Figure 2d**) and BioTransformer results: in lactic acid bacteria proteolytic systems, extracellular proteinases degrade proteins into oligopeptides, and intracellular peptidases further process them into di-and tripeptides enriched in hydrophobic residues (Ter et al., 2024). The pathway enrichment captures the substrate side of this process, while the peptide accumulation represents the product side.

The lipid and non-lipid chemical subclass models provided additional resolution. In cassava, the lipid model showed broad depletion of lysophosphatidylinositols (LPI), lysophosphatidylcholines (LPC), and lysophosphatidylethanolamines (LPE), all 98–100% depleted, along with octadecanoids, and steroids (**Figure 5c**). The non-lipid model captured amino acid and dipeptide, purine ribonucleoside monophosphate, and cyanogenic glycoside depletion, the latter confirming the well-documented detoxification role of cassava fermentation (Montagnac et al., 2009). In cocoyam, the lipid model highlighted N-acyl amines as predominantly depleted, along with sphingomyelins and phosphatidylethanolamine-ceramides, while LPE and oxidized lysophosphatidylserines (O-LPS) showed a mixed response with features both enriched and depleted (**Figure 5d**). The non-lipid model captured amino acid depletion and phenothiazine depletion, though with fewer total hits than cassava, consistent with the more targeted rather than broad metabolic changes in cocoyam (**Figure 5d**).

No model pathways were shared between cassava and cocoyam, reinforcing the tuber-specific nature of the fermentation process. Cassava fermentation consumes purines and carbohydrates, while cocoyam fermentation processes amino acids and generates peptide products.

## 4. Conclusion

Cassava and cocoyam are among the most important staple crops in sub-Saharan Africa, where cassava alone serves as the primary carbohydrate source for millions of people and cocoyam is a widely cultivated secondary staple. For both crops, fermentation is an essential processing step for safe consumption, reducing toxic compounds such as cyanogenic glycosides and calcium oxalate. Despite this dietary importance, both tubers share a starchy composition and similar traditional fermentation practices, and their molecular transformations during fermentation have remained largely unexplored.

Our results show that cassava fermentation is predominantly catabolic, depleting the majority of its detectable chemical space, with the few enriched compounds pointing to microbial biomass rather than new food-relevant metabolites. Cocoyam fermentation both consumed existing metabolites and generated new ones, driven by active proteolytic processing that produced dipeptides and tripeptides not present in the raw tuber. These different responses would not have been captured by proximate composition or targeted assays, and only became apparent through untargeted metabolomics applied comparatively across both tubers. At the same time, the approach captured previously established effects of fermentation in both tubers, such as the depletion of cyanogenic glycosides, confirming that the pipeline recovers known biology alongside novel findings.

Many of the detected compounds are not well represented in existing food or biological databases, indicating that the chemistry of these tubers remained under-characterized. If two tubers that appear nutritionally similar at the proximate level can harbor such different molecular fermentation profiles, the molecular diversity of other traditional fermented African staple foods is likely much larger than currently documented. Understanding these molecular transformations may inform efforts to improve the nutritional quality and safety of fermented tuber products, and to identify relevant bioactive compounds in foods that remain culturally significant but scientifically underexplored.

## Data availability

Raw LC-MS/MS data files are available through the MassIVE data repository (accession no. MSV000101875). Feature-based molecular networking jobs are publicly available on the GNPS2 platform (cassava: https://gnps2.org/status?task=5ee66f5d1e0b415380829dc88d00baf2; cocoyam: https://gnps2.org/status?task=9939fd2e074743238a6020680267357f). MZmine batch files, feature lists, SIRIUS annotation results, and BioTransformer output files are available at https://github.com/bittremieuxlab/cassava_cocoyam_fermentation_study. All data and code needed to reproduce the analyses reported in this study are provided through these repositories.

## Acknowledgements

This research was supported by the Research Foundation – Flanders (FWO G0AGQ24N) and the National Research Foundation (NRF FLA230601112398).

## Author contributions

Alejandro Mendoza Cantu: conceptualization, data curation, formal analysis, investigation, methodology, software, visualization, writing – original draft, writing – review & editing.

Motseoa Mariam Lephatsi: conceptualization, data curation, formal analysis, investigation, methodology, writing – original draft, writing – review & editing.

Yusuf Abass Aleshinloye: investigation, data curation.

Maserufe Faith Phahlane: investigation, formal analysis.

Ntakadzeni Edwin Madala: formal analysis.

Oluwaseun Peter Bamidele: conceptualization, funding acquisition, resources, writing – review & editing.

Ndomele Ndiko Ludidi: conceptualization, funding acquisition, resources, writing – review & editing.

Wout Bittremieux: conceptualization, funding acquisition, methodology, project administration, supervision, writing – review & editing.

Julia M. Gauglitz: conceptualization, funding acquisition, methodology, supervision, writing – review & editing.

Fidele Tugizimana: conceptualization, funding acquisition, methodology, resources, supervision, writing – review & editing.

## Supplementary information

**Supplementary Figure 1.**
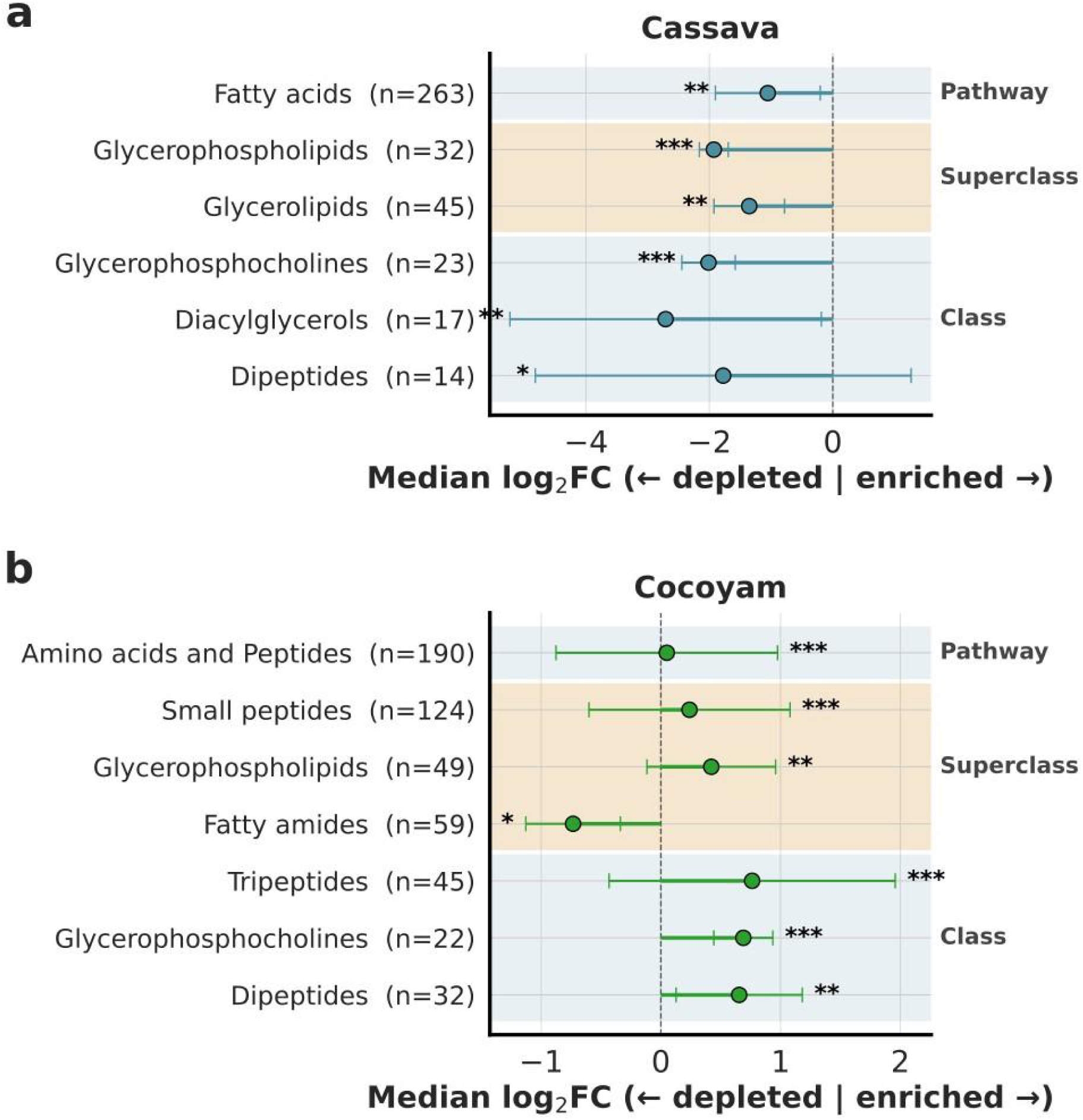
Significant NPClassifier classes across pathway, superclass, and class levels for cassava **(a)** and cocoyam **(b)**. Only classes reaching significance (Benjamini Hochberg FDR adjusted p < 0.05) are shown. Significance was assessed using a Mann-Whitney U test comparing the log₂FC values of features within each class against all other features. Dots represent the median log₂FC across all features assigned to each class, with error bars showing the interquartile range. Class membership sizes are indicated in parentheses. Background shading distinguishes NPClassifier levels: pathway (blue), superclass (orange), class (blue). Significance levels: * p < 0.05, ** p < 0.01, *** p < 0.001.

**Supplementary Figure 2.**
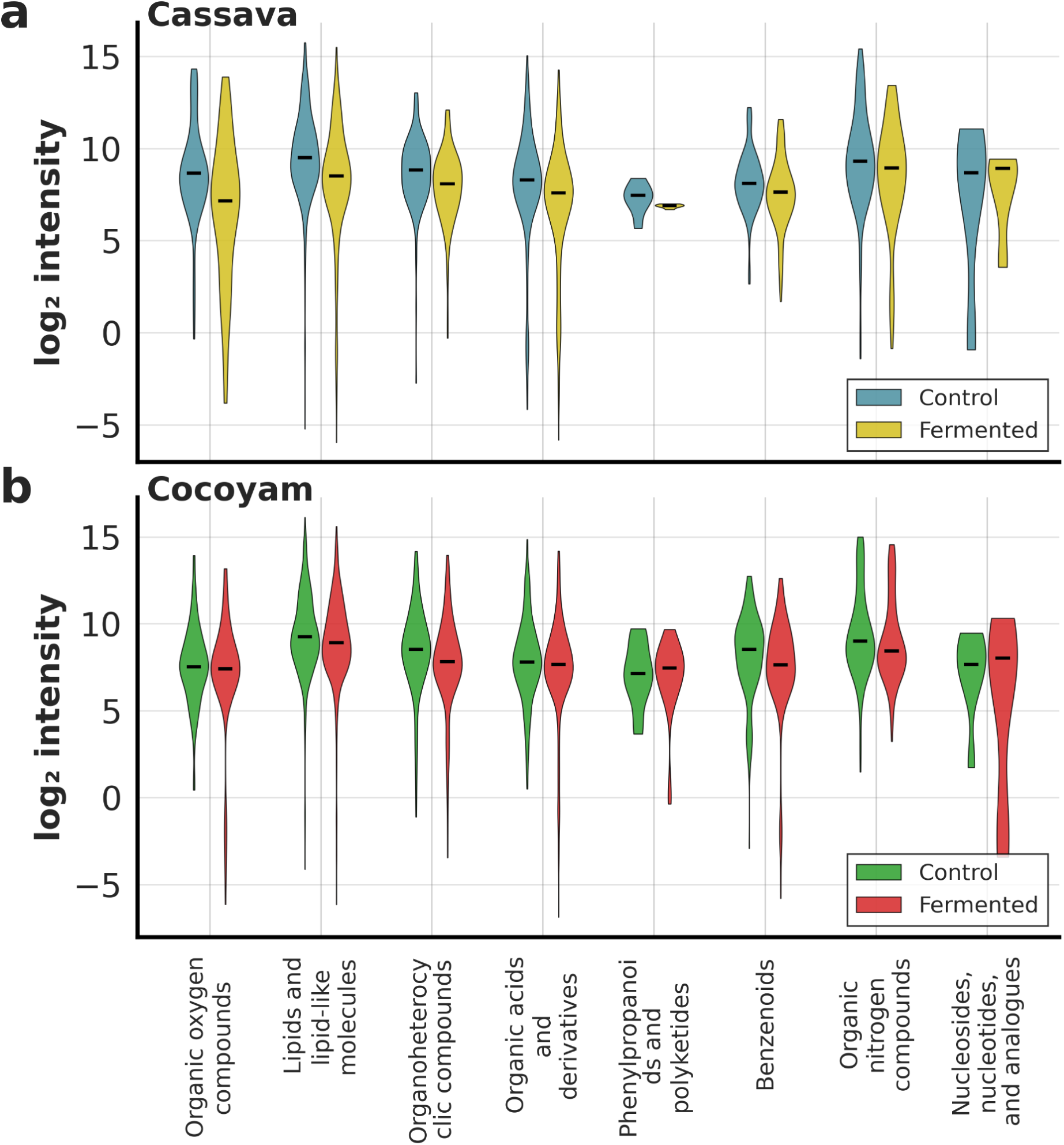
ClassyFire superclass-level intensity distributions for cassava and cocoyam before and after fermentation. Violin plots show log₂-normalized intensity distributions of SIRIUS-annotated features grouped by ClassyFire superclass for **(a)** cassava and **(b)** cocoyam, comparing control and fermented samples. Only features with ClassyFire superclass probability ≥ 0.7 and at least one real measured value in the respective tuber are included.

**Supplementary Figure 3.**
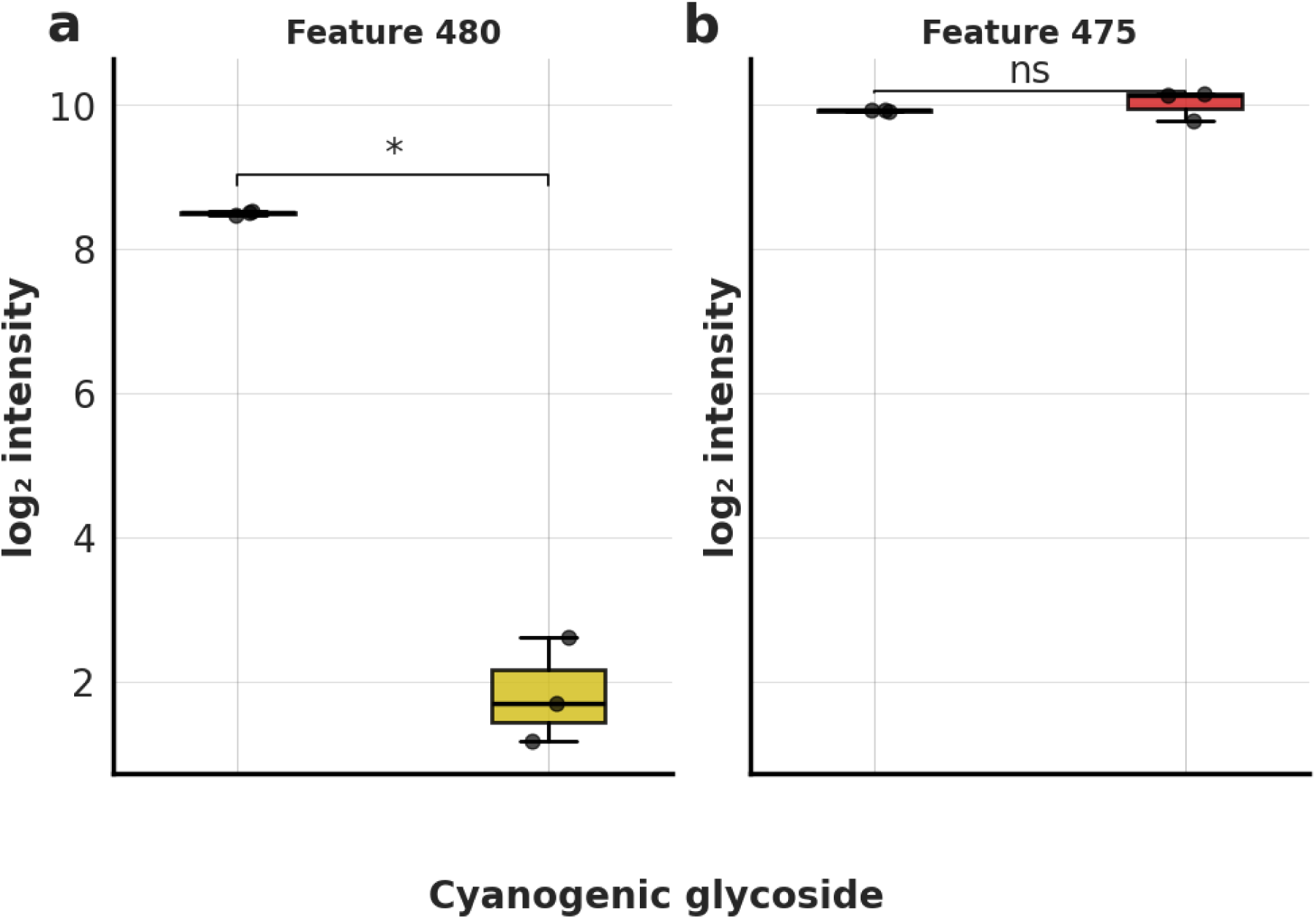
Intensity distribution of a CANOPUS-classified cyanogenic glycoside feature found in both tubers (matching *m*/*z* 490.19 and retention time) in cassava **(a)** and cocoyam **(b)**, comparing control and fermented samples. The feature was significantly depleted after fermentation in cassava (log₂FC = −6.67, Welch’s t test, p < 0.05) but remained unchanged in cocoyam (log₂FC = 0.11, ns).

